# Direct quantitative PCR detects genetic biomarkers of antileishmanial drug resistance in clinical samples from dogs with leishmaniosis

**DOI:** 10.64898/2026.02.03.703521

**Authors:** M. Carrasco-Martin, C. Viñeta, J. Martí-Carreras, X. Roura, L. Ferrer, A. Vila, O. Francino

**Affiliations:** SVGM, Molecular Genetics Veterinary Service, Universitat Autònoma de Barcelona, 08193 Bellaterra, Spain; Hospital Clínic Veterinari, Universitat Autònoma de Barcelona, 08193 Bellaterra, Spain; Nano1Health S.L., Edifici EUREKA, Parc de Recerca, Universitat Autònoma de Barcelona, 08193 Bellaterra, Spain; Departament de Medicina i Cirurgia Animals, Universitat Autònoma de Barcelona, 08193 Bellaterra, Spain; Hospital Veterinario, Universidad Católica de Valencia, 46018 Valencia, Spain

**Keywords:** *Leishmania infantum*, canine, allopurinol, meglumine antimoniate, miltefosine

## Abstract

**Background:** Treatment response in canine leishmaniosis is driven by the dog host, the *Leishmania* parasite, and pharmacological factors, with drug resistance increasingly undermining the effectiveness of therapy. A direct quantitative PCR test (LeishGenR^™^) was applied to 104 clinical samples from 95 dogs in the Mediterranean area diagnosed with leishmaniosis in veterinary clinical settings and testing positive for *Leishmania infantum* by PCR. The assay enabled rapid detection of genetic drug-resistance biomarkers for allopurinol (*metk*), meglumine antimoniate (*mrpa*), and miltefosine (*LdMT*), providing a clinically relevant, timely alternative to culture-based approaches by directly analyzing circulating *Leishmania infantum* amastigotes.

**Results:** The assay (LeishGenR^™^) showed high specificity (100%) and sensitivity (>87.5%) for genetic drug-resistance profile assignment and a strong correlation with whole-genome sequencing for gene copy number assessment (*metk*: r = 0.878; *mrpa*: r = 0.943 and *LdMT* = 0.691). Genetic drug-resistance biomarkers were detected in 24.3% of *L. infantum* DNA from clinical samples analyzed (20/82; 95% CI 16.3-34.6)), most commonly for allopurinol (13.4%; 95% CI 7.6-22.4), then meglumine antimoniate (9.4%; 95% CI 4.6-18.2), and for miltefosine (5.4%; 95% CI 1.8-14.8). Prevalence was higher in dogs previously treated for leishmaniosis.

**Conclusion:** This study demonstrates the ability to detect genetic biomarkers of drug resistance in *L. infantum* directly from clinical samples of dogs with leishmaniosis. This method enables rapid, precise detection of genomic biomarkers, circumventing delays associated with culture-based methods and supporting more effective clinical management and surveillance. Among dogs with high parasitemia referred to clinics in Mediterranean regions sampled in this study, the findings reveal a significant prevalence of circulating *L. infantum* strains carrying genomic drug resistance biomarkers to standard treatments for canine leishmaniosis.

## 1. Background

Canine leishmaniosis (CanL) is a vector-borne zoonotic disease caused by the parasite *Leishmania infantum*. The dog is the main companion animal reservoir for *L. infantum*, the primary etiological agent of human leishmaniosis in the Mediterranean area. Dogs are also the domestic animal most severely affected clinically by this disease [1,2]. Previous studies using PCR-based assays revealed a significant prevalence of subclinical infections [3], highlighting that a substantial proportion of infected dogs remain without clinical signs, serving as potential reservoirs and exhibiting infectivity to the main vector in the Mediterranean region, *Phlebotomus pernicious* [4,5].

The CanL is considered endemic in the Mediterranean basin and is becoming a threat in central and northern European countries [6–8]. The CanL poses a significant public health concern in Western Mediterranean countries, where approximately 2.5 million dogs are infected with the parasite *L. infantum* [9].

The most recommended and effective treatment protocols for CanL are a combination of antileishmanial drugs, usually, one month of meglumine antimoniate or miltefosine, together with at least 6 months of allopurinol [1,10]. These protocols may induce temporary or permanent remission but do not entirely result in complete parasitological cure. Relapses of CanL following treatment are frequent [11,12]. Treatment efficacy in CanL is influenced by drug-related factors; host-related factors, such as the immune response or the presence of other comorbidities; and parasite-specific factors, with drug resistance emerging as a critical factor undermining treatment effectiveness [13–15]. Long-term or overused antileishmanial treatment likely drives drug resistance by applying selective pressure on dog parasites [14–19]. Drug resistance is of significant concern primarily in veterinary health care, but additionally for human health as some drugs are also used to treat leishmaniosis in humans [13].

*Leishmania infantum* isolates resistant to allopurinol [17,20], meglumine antimoniate [21], and miltefosine [22] have been reported in clinical samples from dogs with leishmaniosis. Copy number variation (CNV) on different *L. infantum* genes have been reported as potential genetic drug-resistance biomarkers (GDRB) for allopurinol, meglumine antimoniate, and miltefosine, such as *metk* [23], multidrug-resistance protein A gene (*mrpa)* [24], and *L. donovani* miltefosine transporter gene (*LdMT)* [25,26], respectively. A previous study [27] examined the prevalence of genetic drug-resistance biomarkers (GDRB) in cultured isolates of *L. infantum* from the Mediterranean region. The most frequently identified GDRB in canine isolates were allopurinol and meglumine antimoniate, with prevalence rates ranging from 22.7% to 50% [27]. However, these results are derived from cultured isolates and may be susceptible to genomic alterations that occur during *Leishmania* culture adaptation [19]. The prevalence of GDRB in clinical samples from dogs with leishmaniosis in endemic areas remains largely unknown.

The main objective of this study was to assess the ability of a direct quantitative PCR (qPCR) testing-based method to detect genetic biomarkers of drug resistance in *L. infantum* from clinical samples of dogs with leishmaniosis. Utilizing clinical samples rather than culture-based methods allows for quicker results, facilitating timely treatment adjustments, clarifying reasons for treatment failure, and differentiating between drug resistance and an inadequate immune response. The second objective was to analyze clinical samples from dogs diagnosed with leishmaniosis and to describe the prevalence of GDRB in relation to allopurinol, meglumine antimoniate, and miltefosine, with the aim of understanding the status of circulating *L. infantum* strains in Mediterranean dogs with leishmaniosis.

## 2. Material and methods

### 2.1. Study design and sample collection

The present study includes 104 samples from 95 dogs diagnosed with leishmaniosis in veterinary clinical settings and tested positive for *Leishmania infantum* by PCR. Samples were collected from 2018 to 2025 from different veterinary clinical settings in Spain (Andalucia (7), Aragón (1), Balearic Islands (3), Catalonia (63), Euskadi (1), Extremadura (2), Madrid (5), Murcia (1), Region of Valencia (7)), and Greece (4). Additional samples (10) came from southern Europe, though their exact origin could not be determined. Clinical samples were obtained from six biological sites: peripheral blood (n = 58), lymph node (n = 30) and bone marrow aspirates (n = 12), skin biopsy (n = 2), and one each from joint and pericardium effusions. A brief, structured questionnaire was completed by the veterinary practitioners, recording the dog’s age, sex, clinical signs suggestive of leishmaniosis, and prior antileishmanial treatment history.

### 2.2. DNA extraction and qPCR for Leishmania detection

DNA was obtained from 0.1 mL of lymph node or bone marrow aspirates, or from 0.5 mL of peripheral whole blood or other fluids, as previously described [28]. Briefly, samples were transferred to a 1.5-ml sterile tube and washed with TE Buffer (pH 8.0) to disrupt the erythrocyte membrane until the leukocyte pellet became whitish. Then, leukocyte cells were lysed by adding 0.1 ml of a mixture of K-Buffer (50 mM KCl, 10 mM TRIS, pH 8.0, 0.5% Tween-20) and proteinase K (20mg/ml), previously prepared, and incubated at 56ºC for at least 3-4 hours. Once digestion was complete, proteinase K was inactivated at 90 °C for 10 minutes, and DNA extraction was ready for downstream analysis.

For skin samples, DNA extraction was performed using the PureLink Genomic DNA Mini Kit (Invitrogen, Carlsbad, CA, USA) according to the manufacturer’s recommendations.

*Leishmania* detection in the extracted DNA was confirmed using qPCR with primers and a TaqMan-MGB probe designed to target conserved DNA regions of the kinetoplast minicircle DNA from *L. infantum*, as previously described [29].

### 2.3. Leishmania GDRB testing

DNA from the clinical samples was subsequently tested with the molecular test LeishGenR^™^ (Nano1Health SL, Barcelona, Spain), a direct qPCR-based method for detecting GDRB in *L. infantum*. The LeishGenR^™^ TaqMan-MGB probes and primers target specific regions of *L. infantum* genome to assess CNV in *metk, mrpa*, and *LdMT* genes, reported as potential GDRB for allopurinol [23], meglumine antimoniate [24], and miltefosine [25,26], respectively, as well as a DNA region of the housekeeping control gene, meant to be stable in two copies and used for CNV determination by the delta Ct method. Each amplification run included both positive and negative controls.

### 2.4. Assessment of efficiency, sensitivity, and specificity of the LeishGenR^™^ assay

Three replicates of 10-fold serial dilutions of parasite DNA from a culture of *L. infantum* (LCAN/ES/2023/CATB109) and a synthetic DNA construct containing a single copy of the four amplicon sequences were used to evaluate the LeishGenR^™^ test’s efficiency via the TaqMan-MGB housekeeping gene assay. Sensitivity and specificity for GDRB determination were assessed using leftover DNA from cultured *L. infantum* isolates. The copy numbers of *metk, mrpa*, and *LdMT*, together with the genomic drug resistance biomarker profile from whole-genome sequencing (WGS)[27], served as the gold standard for evaluating LeishGenR^™^ performance.

### 2.5. Statistical analysis

The repeatability of LeishGenR^™^ in assessing antileishmanial GDRB profiles was determined by analyzing two *L. infantum* isolates (S1: MCAN/IT/2020/1265; S2: MCAN/IT/2022/2885) in 12 independent replicates per biomarker. Data normality distribution was verified using the Shapiro–Wilk test. A t-test was performed to estimate the population’s mean CNV and its 95% confidence interval, to assess whether the estimated CNV surpassed the predefined biomarker detection thresholds.

Wilson’s method was used to calculate 95% confidence intervals (95% CIs) for the prevalence of GDRB.

For descriptive treatment history analysis, a dog was considered as treated if the drug had been administered at some point in its clinical history.

## 3. Results

### 3.1. Verification of the LeishGenR^™^ assay performance for GDRB detection

The drug-resistance biomarker assay LeishGenR^™^ (Nano1Health SL) is a direct qPCR-based method for assessing CNV as GDRB in *L. infantum*. It targets the *metk* (allopurinol), *mrpa* (meglumine antimoniate), and *LdMT* (miltefosine) genes, using a stable two-copy housekeeping control gene for CNV normalization.

The qPCR TaqMan assay targeting the two-copy control gene was used to determine LeishGenR^™^ efficiency. Standard curves for both synthetic and *L. infantum* isolate DNA showed high and similar amplification efficiency. The qPCR efficiency for synthetic DNA reached 99.55%, and for the cultured isolate 97.17%. The dynamic range for gene CNV calling was set to 6 logs, and the limit for GDRB analysis was 10 parasites (20 genome copies) per qPCR reaction, with a correlation of 0.99 and an efficiency of 99.44%.

Prior to clinical sample testing, *L. infantum* cultured isolates that had been previously analyzed by WGS were tested [27,30]. The WGS data is considered the gold standard for determining true positives and true negatives in the performance analysis. A total of 22 isolates for *metk* and *mrpa*, and 16 *LdMT* were tested.

On cultured isolates, LeishGenR^™^ showed 100% specificity for *metk, mrpa*, and *LdMT*. Sensitivity was 87.5% for *metk* and 100% for *mrpa*. Agreement in genomic-response profile assignment between LeishGenR^™^ and WGS was nearly perfect (Cohen’s Kappa: 0.81 for *metk*, and 1 for *mrpa*; Table 1). In addition, Pearson’s correlation showed strong agreement for CNV (*metk*: r = 0.878, p < 0.001; *mrpa*: r = 0.94, p < 0.001; *LdMT*: r=0.691, p<0.001) (Figure 1; Table 1). Additional file 1 shows CNV values obtained through WGS and qPCR (see Additional file 1).

**Table 1.** Summary of LeishGenR^™^ test metrics performance.

|  | *metk* CNV profile<br>(Allopurinol) | *mrpa* CNV profile<br>(Meglumine antimoniate) | *LdMT* CNV profile<br>(Miltefosine) |
| --- | --- | --- | --- |
| **Profile assessment performance** |  |  |  |
| Sensitivity (%) | 87.5 | 100 | NA* |
| Specificity (%) | 100 | 100 | 100 |
| Profile agreement (k, P) | 0.81, < 0.001 | 1, < 0.001 | NA* |
| **CNV assessment performance** |  |  |  |
| Pearson coefficient (r, P) | 0.914, < 0.001 | 0.943, < 0.001 | NA* |
| **Repeatability** |  |  |  |
| S1: mean (IC 95%) | 2.74 (2.57, 2.91)<br>df = 11 | 4.11 (3.94, 4.29)<br>df = 11 | 2.2 (2.00, 2.39)<br>df = 7 |
| S2: mean (IC 95%) | 2.64 (2.48, 2.79)<br>df = 11 | 2.06 (1.86, 2.27)<br>df = 11 | 1.83 (1.71, 1.95)<br>df = 11 |

NA\*: The assessment of sensitivity for *LdMT* was not meaningful because all tested cultures were negative for this biomarker by WGS.

**Figure 1:**
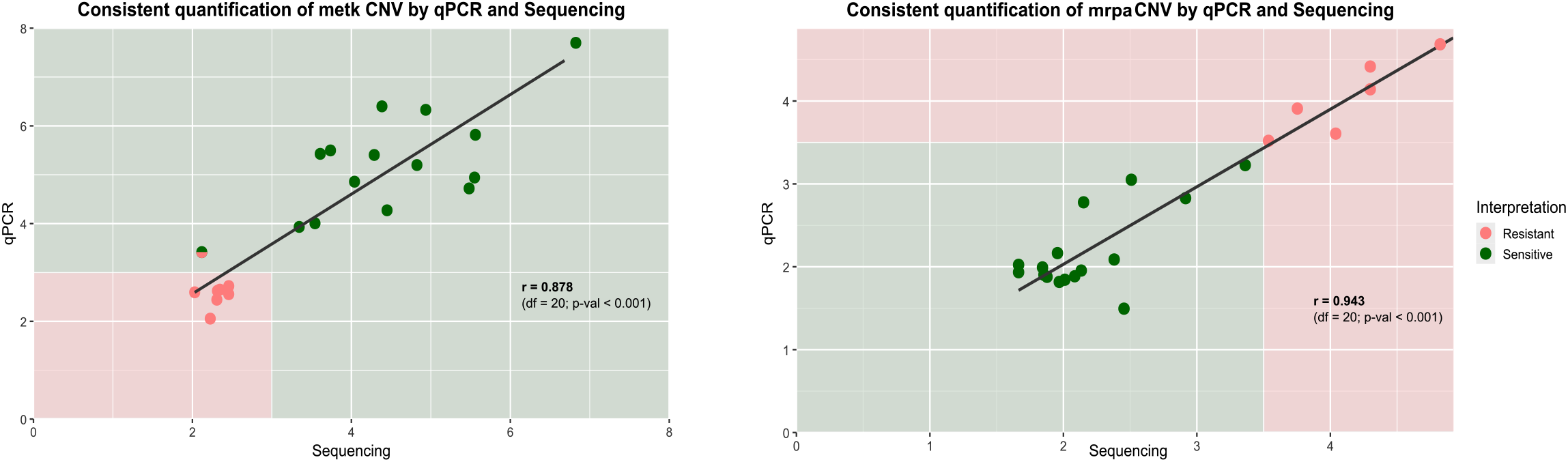
Correlation between LeishGenR^™^ and WGS for *metk* (left) and *mrpa* (right) CNV assessment. Scatter plot of *metk* and *mrpa* genes CNV for the two techniques. Red: resistant. Green: sensitive.

The assessment of *LdMT* sensitivity was not meaningful because all tested cultures were negative for this biomarker by WGS. The validation of the *LdMT* assay was completed by confirming the consistent detection of *LdMT* and the control gene copies in the DNA synthetic construct in the qPCR reaction.

Twelve replicates of two *L. infantum* cultures were analyzed using LeishGenR^™^ to assess the repeatability of the GDRB assay. The assay consistently classified sample 1 (S1) as resistant to both allopurinol (mean *metk* CN: 2.75 (95% CI 2.57–2.91, df = 11) and meglumine antimoniate (mean *mrpa* CN: 4.11 (95% CI 3.94–4.29, df = 11) and sensitive to miltefosine (mean *LdMT* CN: 2.2 (95% CI 2.00–2.39, df = 11). Sample 2 (S2) was classified as resistant to allopurinol (mean *metk* CN: 2.64 (95% CI 2.48–2.79, df = 11), and sensitive to both meglumine antimoniate (mean *mrpa* CN: 2.06 (95% CI 1.86–2.27, df = 11) and miltefosine (mean *LdMT* CN: 1.83 (95% CI 1.71– 1.95, df = 11).

### 3.2. GDRB prevalence for allopurinol, meglumine antimoniate and miltefosine

A total of 104 clinical samples from dogs with leishmaniosis were collected, including 58 peripheral whole-blood samples, 30 lymph node and 12 bone marrow aspirates, 2 skin biopsies, and one each of joint and pericardium effusions. All samples tested positive for *Leishmania infantum* qPCR [29].

Eighty-two of the 104 samples (79%) met the criteria for GDRB analysis using LeishGenR^™^, with sufficient parasite load (≥10 parasites per qPCR reaction). Bone marrow and lymph node aspirates had higher success rates for GDRB analysis (lymph node: 90%, 27/30; bone marrow: 83%, 10/12) than peripheral blood samples (71%, 41/58). All 82 suitable samples were assessed for allopurinol GDRB. Of these, 74 were also tested for meglumine antimoniate GDRB, and 55 samples with leftover DNA were analyzed for all three GDRB (allopurinol, meglumine antimoniate, and miltefosine).

Twenty-two samples had insufficient parasite load (<10 parasites per qPCR reaction) for accurate CNV analysis for GDRB assessment by qPCR. These included 17 peripheral blood, 3 lymph nodes and 2 bone marrow aspirates. Sample metadata and a summary of GDRB results are provided in Supplementary Material 1.

Table 2 shows the prevalence of the three GDRB in the clinical samples analyzed. At least one antileishmanial GDRB was detected in 24.3% (20/82; 95% CI 16.3-34.6) of the samples tested. Prevalence for allopurinol GDRB was 13.4% (95% CI 7.6-22.4), for meglumine antimoniate 9.4% (95% CI 4.6-18.2), and for miltefosine 5.4% (95% CI 1.8-14.8) (Figure 2).

**Table 2.**
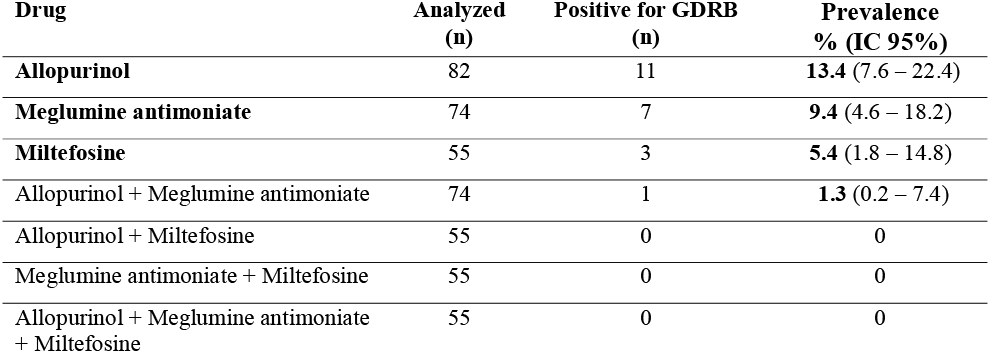
Prevalence of genetic drug-resistance biomarkers (GDRB) in *L. infantum* from clinical samples of dogs with leishmaniosis.

| Drug | Analyzed (n) | Positive for GDRB (n) | Prevalence % (IC 95%) |
| --- | --- | --- | --- |
| **Allopurinol** | 82 | 11 | **13.4** (7.6 – 22.4) |
| **Meglumine antimoniate** | 74 | 7 | **9.4** (4.6 – 18.2) |
| **Miltefosine** | 55 | 3 | **5.4** (1.8 – 14.8) |
| Allopurinol + Meglumine antimoniate | 74 | 1 | **1.3** (0.2 – 7.4) |
| Allopurinol + Miltefosine | 55 | 0 | 0 |
| Meglumine antimoniate + Miltefosine | 55 | 0 | 0 |
| Allopurinol + Meglumine antimoniate + Miltefosine | 55 | 0 | 0 |

**Figure 2:**
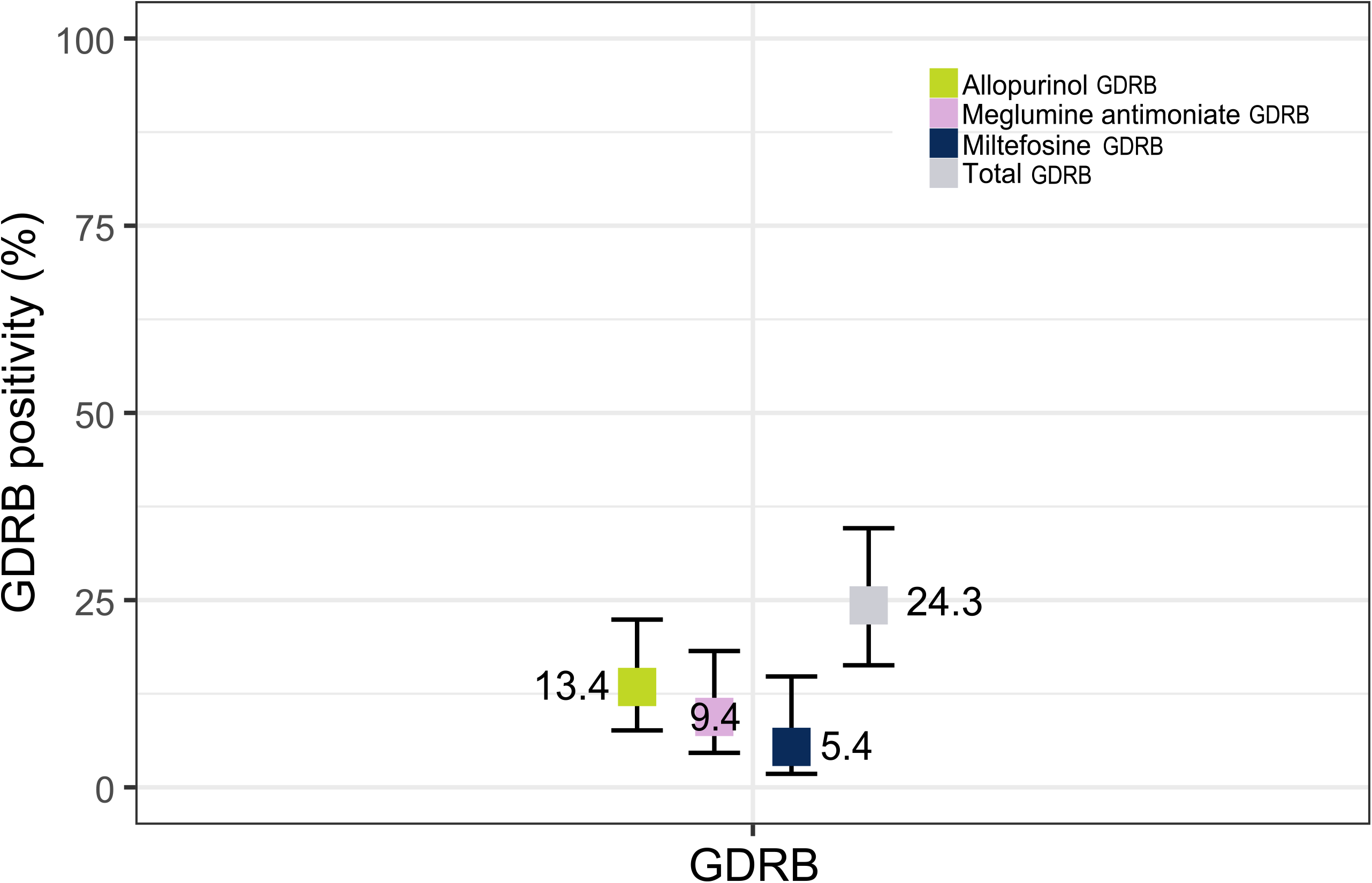
Detection of at least one GDRB in 24.3% of DNAs extracted from L. infantum in clinical samples. GDRB positivity rates (GDRB positivity rate ± CI 95%) for allopurinol (green), meglumine antimoniate (purple), miltefosine (dark blue), and the combination of all three drugs (Total; gray).

There were no statistically significant differences in the prevalence of GDRB positivity among the three drugs (Fisher’s Exact Test; P = 0.316). Most positive cases involved monoresistance, with only one sample (1.3%; 95% CI 0.2-7.4) testing positive for two antileishmanial GDRB: allopurinol and meglumine antimoniate. No sample showed combined allopurinol-miltefosine or meglumine antimoniate-miltefosine GDRB.

### 3.2. Factors associated with GDRB detection

Clinical histories for 34 dogs, corresponding to 37 clinical samples, were gathered. Most dogs (94%, 32/34) had relapsing clinical leishmaniosis, and 6% (2/34) were diagnosed with CanL for the first time. Dogs commonly presented dermatological lesions (8), anemia (5), proteinuria (5), and azotemia and epistaxis (3) at the time of sample isolation for GDRB testing. More details on clinical signs can be found in Additional file 2.

Regarding treatment, at the time of sample isolation, the dog received allopurinol in 33 of 37 samples (90%), meglumine antimoniate in 24 of 37 (65%), and miltefosine in 11 of 37 (29%) (Figure3a).

To study the potential association between GDRB detection and treatment history, the exploratory analysis focused on the samples tested for each GDRB and with treatment data available: 34 out of the 82 samples analyzed for allopurinol biomarker, 32 out of the 74 tested for meglumine antimoniate one, and 25 out of the 55 analyzed for miltefosine one (Table 3).

**Table 3.** Relation of total samples analyzed for GDRB and clinical history.

| Drug | Total analyzed<br>for GDRB | Clinical history | Treated | Non-treated |
| --- | --- | --- | --- | --- |
| Allopurinol | 82 | 34/82 | 31/34 | 3/34 |
| Meglumine<br>antimoniate | 74 | 32/74 | 21/32 | 11/32 |
| Miltefosine | 55 | 25/55 | 8/25 | 17/25 |

In this exploratory dataset with clinical history and previous treatment data available, treated dogs showed higher observed GDRB prevalences than untreated ones (Figure 3b). For allopurinol, the *metk* biomarker was identified in 11.8% (4/34) of the exploratory dataset, corresponding to 12.9% (4/31) in previously treated dogs, and was not detected in untreated dogs (0/3). For meglumine antimoniate, the overall *mrpa* biomarker prevalence was 12.5% (4/32), rising to 19% for samples from meglumine antimoniate-treated dogs (4/21), while no meglumine antimoniate resistance biomarker was detected in non-treated dogs (0/11). Finally, the overall prevalence of the miltefosine *LdMT* biomarker was 8% (2/25), with 12.5% in samples from treated dogs (1/8) and 5.9% in samples from non-miltefosine treated dogs (1/17) (Figure 3b).

**Figure 3:**
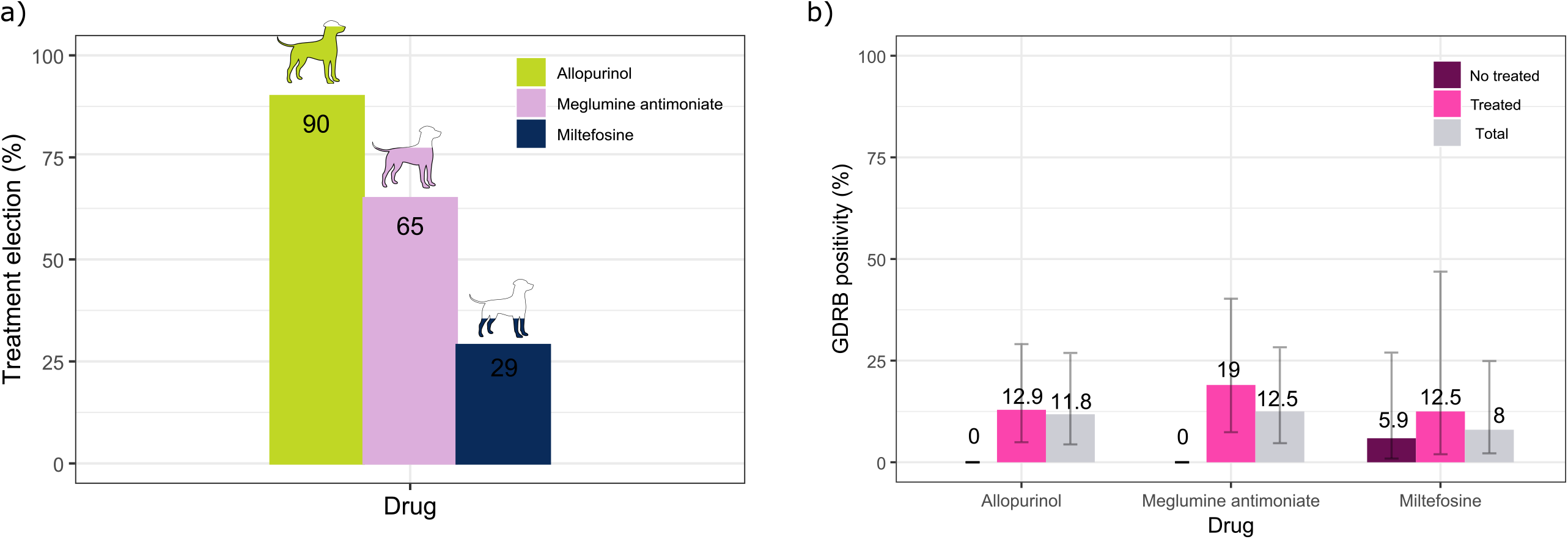
Antileishmanial GDRB detection in treated dogs for the three drugs used. Bar plot showing percentage of samples treated with allopurinol (green), meglumine antimoniate (purple), and miltefosine (blue) **(a)**. Overall prevalence of antileishmanial GDRB in the clinical samples analyzed (Total; gray), compared with treated dogs (Treated; pink) and non-treated dogs (No Treated; purple) for allopurinol, meglumine antimoniate, and miltefosine. Error bars showing 95% CI. **(b)**.

## 4. Discussion

This study shows the ability of the LeishGenR^™^ qPCR-based assay to detect genetic biomarkers linked to drug resistance in *L. infantum* for allopurinol, meglumine antimoniate, and miltefosine directly from clinical samples of dogs with leishmaniosis.

The assay achieves high qPCR efficiency for the endogenous control gene (99.55% with a synthetic DNA construct and 97.17% with an *L. infantum* culture isolate) both exceeding the threshold for well-amplified standards [31]. The use of a synthetic DNA standard as control further streamlines workflows, reduces costs, and ensures robust, reproducible results. Standard curve results demonstrated that this test is suitable for CNV estimation in samples with more than 10 parasites (20 genome copies) in the qPCR reaction, corresponding to the lowest concentration at which the assay shows ≤35% coefficient of variation [32]. Comparable numbers of genome copies (10–100 per reaction) have been reported in previous studies for CNV discrimination by qPCR [32,33]. Therefore, it is important to note that this method requires sufficient parasite load, limiting its applicability to clinical samples with high parasite concentrations. As a future development, testing the method in digital PCR to enhance CNV estimation accuracy for low-parasitemia samples could be of interest, whereas qPCR-based CNV calls remain robust for samples with higher parasitemia.

To validate the method’s reliability in GDRB prediction, the GDRB profiles of *L. infantum* culture isolates were compared with those obtained by WGS, which was considered the gold standard. Correlation between WGS and qPCR copy number in *metk* (allopurinol GDRB) and *mrpa* (meglumine antimoniate GDRB) in the *L. infantum* isolates had a Pearson coefficient above 0.9 (P < 0.001). Similarly, a previous study on the chicken genome WGS validated a subset of CNVs using qPCR, demonstrating comparable concordance between WGS and qPCR measurements, as evidenced by high qPCR validation rates [34]. Overall, no differences in GDRB profile assignment were found by qPCR or WGS, except for one sample (positive for allopurinol GDRB by WGS and negative by qPCR), yielding a false-negative rate of 12.5% for metk. Similarly, a 22.43% false negative rate was reported in the above-referenced chicken-genome CNV study [34].

Taken together, the present results show that LeishGenR^™^ enables rapid, accurate detection of *L. infantum* GDRBs from direct analysis of clinical samples, expediting clinical decision-making and avoiding the need for parasite *in vitro* expansion, which can cause genomic alterations [19], and providing significant advantages over traditional culture-based analysis. The methodological advancements demonstrated in this study lay the groundwork for broader implementation of molecular diagnostics in GDRB.

This study also demonstrated the presence of clinical strains of *L. infantum* harboring GDRB to allopurinol, meglumine antimoniate, and miltefosine circulating among dogs with leishmaniosis in the Mediterranean basin. In this cohort, 24.3% of *L. infantum* DNA samples from clinical specimens tested positive for at least one antileishmanial GDRB, with allopurinol GDRB being the most prevalent (13.4%), followed by meglumine antimoniate (9.4%) and miltefosine (5.4%).

These findings contrasted with previous studies on cultured *L. infantum* isolates from both dogs and humans in the same region, which reported a higher overall prevalence of GDRB for these three drugs (48.5%; 95% CI 32.9-64.4)[27]. This previous study also observed the same pattern in the prevalence order for the three antileishmanial GDRB studied in cultured dog isolates[27]. However, current clinical samples showed lower prevalence rates than those from cultured isolates from dogs. The differences may reflect factors such as geographical origin, previous treatment history, the complexity of the leishmaniosis cases sent to the reference institutions, or potential culturing effects in high passage cultures [19].

Among the resistance biomarkers analyzed, allopurinol GDRB was the most frequent (13.4%). This reflects its role as the primary leishmaniostatic drug for long-term management of canine leishmaniosis[1,10–12]. All allopurinol GDRB-positive samples were from dogs previously treated with this drug. This supports the association between drug exposure and resistance development [18]. There is a potential link between allopurinol resistance and disease relapse [17]. Further longitudinal studies are needed to clarify this relationship, particularly in dogs with complex clinical histories. Alternative strategies for controlling parasite load, such as immunotherapy, may also be warranted for effective parasite control in cases of resistance[11,12,35].

Among leishmanicidal drugs, meglumine antimoniate GDRB was the most frequently detected (9.4% of samples), in line with its administration to 65% of treated dogs. The persistence of resistance through sand fly transmission and potential fitness advantages of resistant strains are concerning both a clinical and epidemiological perspective [36]. Miltefosine resistance was less frequent (5.4%), consistent with its more recent and less widespread use (32% of cases) and the reduced fitness of resistant parasites in mammalian hosts, which may limit their spread despite the maintenance of resistance through the vector [37], in addition to being a much more recent leishmanicidal treatment option in CanL management [11]. Importantly, these prevalence estimates should be interpreted with caution due to limited and uneven sample sizes, which may introduce bias and challenge their generalizability. Nevertheless, understanding resistance patterns to these drugs in dogs is critical from a One Health perspective, as both remain key components of human leishmaniosis therapy.

Co-occurrence of multiple drug resistance biomarkers was rare (1.3%), supporting current recommendations for combination therapy with drugs that have distinct mechanisms to delay resistance [38,39]. However, vigilance remains necessary, as resistance potential persists and could impact canine and human health. Overuse of allopurinol in dogs with leishmaniosis may lead to drug resistance, threatening the health of both species, even though allopurinol is not part of the therapeutic arsenal for leishmaniosis treatment in humans.

Finally, the clinical application of GDRB detection could improve the management of CanL, especially in endemic regions. Clinical outcomes are multifactorial and depend on factors like drug resistance, treatment compliance, inadequate immune response, and other comorbidities [1,10,12]. Therefore, longitudinal follow-up of dogs with leishmaniosis is essential to assess the consistency of clinical relapse and resistance to antileishmanial agents, as also described in previous works [20]. The next phase of research should involve longitudinal clinical monitoring of dogs (collection of hematological, biochemical, and diagnostic outcomes, specifically qPCR and GDRB results) to determine the consistency of clinical relapse and genomic resistance to the antileishmanial drugs. Further, early identification of resistance may help avoid unnecessary or ineffective treatments, reduce the risk of relapse, and distinguish treatment failure due to drug resistance versus inadequate immune response [14,15]. Furthermore, since vectors can also transmit resistant parasites, integrating GDRB screening into clinical practice has important implications not only for canine health but also for public health within a One Health framework.

## 5. Conclusion

The present study demonstrated the ability of direct qPCR testing of clinical samples from dogs with leishmaniosis to detect genetic biomarkers of antileishmanial resistance. Direct clinical sample testing enables rapid and accurate detection of GDRB, bypassing the delays of culture-based methods and supporting more effective clinical management and surveillance, potentially reducing relapses and the emergence of drug-resistant subpopulations. Furthermore, this study suggests that the non-negligible percentage of circulating *L. infantum* strains positive for GDRB should be a cause for concern. Given the interest of One Health approach to leishmaniosis and the risk of transmission of resistant *L. infantum* strains, ongoing surveillance and further studies—including larger, multicenter investigations—are warranted to monitor resistance trends and guide both veterinary and human health strategies.

## Declarations

### Ethics approval and consent to participate

Not applicable

### Consent for publication

Not applicable

### Availability of data and materials

The datasets supporting the findings of this article are included within the publication.

### Authors’ contributions

MCM, OF, and XR conceived and designed the experiments and conducted the formal analysis. MCM performed the laboratory experiments and analyzed and interpreted the data from clinical samples. CV and AV collected part of the clinical samples and recorded the dog’s clinical history. MCM and OF wrote the original draft. OF, JMC, XR, and LF reviewed the manuscript. All authors read and approved the final version of the manuscript.

## Acknowledgements

The authors would like to thank all the veterinarians for their efforts in collecting the samples.

